# SUMO E3 ligase Mms21 prevents spontaneous DNA damage induced genome rearrangements

**DOI:** 10.1101/141028

**Authors:** Jason Liang, Bin-zhong Li, Alexander P. Tan, Richard D. Kolodner, Christopher D. Putnam, Huilin Zhou

## Abstract

Mms21, a subunit of the Smc5/6 complex, possesses an E3 ligase activity for the Small Ubiquitin-like MOdifier (SUMO), which has a major, but poorly understood role in genome maintenance. Here we show mutations that inactivate the E3 ligase activity of Mms21 cause Rad52- and Pol32-dependent break-induced replication (BIR), which specifically requires the Rrm3 DNA helicase. Interestingly, mutations affecting both Mms21 and the Sgs1 helicase, but not sumoylation of Sgs1, cause further accumulation of genome rearrangements, indicating the distinct roles of Mms21 and Sgs1 in suppressing genome rearrangements. Whole genome sequencing further revealed that the Mre11 endonuclease prevents microhomology-mediated translocations and hairpin-mediated inverted duplications in the *mms21* mutant. Consistent with the accumulation of endogenous DNA lesions, *mms21* cells accumulate spontaneous Ddc2 foci and display a hyper-activated DNA damage checkpoint. Together, these findings support a new paradigm that Mms21 prevents the accumulation of spontaneous DNA lesions that cause diverse genome rearrangements.

## Introduction

The <u>S</u>mall <u>U</u>biquitin-like <u>MO</u>difier (SUMO) regulates many biological processes through its covalent attachment to lysine residues on target proteins via a cascade of an E1-activating enzyme (Aos1-Uba2 in *Saccharomyces cerevisiae*), an E2-conjugating enzyme Ubc9, and one of several SUMO E3 ligases [1]. Three mitotic SUMO E3 ligases (Siz1, Siz2 and Mms21/Nse2) have been identified in *S. cerevisiae*, and these enzymes control substrate-specific sumoylation *in vivo*. Siz1 and Siz2, two paralogs of the PIAS family SUMO E3 ligases [2], catalyze the bulk of intracellular sumoylation [3,4], while the SUMO E3 ligase Mms21 has fewer known substrates [5,6]. This mitotic SUMO pathway is essential for cell viability in *S. cerevisiae*; deletions of *AOS1, UBA2*, or *UBC9* cause lethality, as does combined inactivation of all three mitotic SUMO E3 ligases [5]. In contrast, sumoylation of proteins by Mms21 is not necessary for viability in the presence of Siz1 and Siz2 in *S. cerevisiae* nor do mice require the SUMO E3 ligase activity of the mouse Mms21 ortholog NSMCE2 [7], indicating some redundancy between mitotic E3 ligases.

Mms21 is an integral subunit of the Smc5/6 complex [8]. The Smc5/6 complex belongs to the evolutionarily conserved SMC family proteins and acts in maintaining chromosome integrity [9]. Loss of the Mms21 SUMO E3 ligase activity causes aberrant increases in homologous recombination (HR) intermediates and accumulations of gross chromosomal rearrangements (GCRs) in *S. cerevisiae* [4,10–12]. Consistent with this, mutations in human *NSMCE2/MMS21* cause increased sister chromatid exchange (SCE) [13] and have been recently linked to DNA replication and/or repair defects and primordial dwarfism [14].

How sumoylation by Mms21 acts to suppress the accumulation of HR intermediates and GCRs is not known. These phenotypes could be attributed to a failure in resolving recombination intermediates and/or an elevated incidence of DNA lesions that are repaired by HR. These phenotypes, however, are reminiscent of those found in cells lacking the Sgs1 helicase [10,15]. Sgs1, the *S. cerevisiae* ortholog of the human BLM helicase that is deficient in patients with Bloom syndrome, has well-documented roles in resolving HR intermediates as well as participating in resection of DNA double strand breaks (DSBs) [16–18]. The similarity between the phenotypes caused by *sgs1Δ* and *mms21* E3 ligase-defective mutations raises the possibility that Mms21 and Sgs1 might function together to regulate or prevent HR [10]. In support of this model, two recent studies showed that sumoylation of Sgs1/BLM by Mms21/NSMCE2 prevents the accumulation of aberrant HR intermediates induced by DNA alkylation damage [19,20].

In contrast, several lines of evidence suggest that Mms21 and Sgs1 act in separate pathways to maintain genome stability. The *sgs1* mutations that eliminate DNA damage-induced sumoylation of Sgs1 by Mms21 do not cause appreciable sensitivity to DNA damaging agents [19,20], unlike that seen for *mms21* E3 ligase defective mutants and *sgs1Δ* mutants [5,10]. Similarly, combining mutations affecting *NSMCE2/MMS21* and *BLM/SGS1* caused synthetic growth defects and increased SCE in mouse B cells [7]. We previously demonstrated that Esc2, a protein containing two SUMO-like domains with an important role in genome maintenance [11,21], functions together with Mms21 in controlling intracellular sumoylation and suppressing GCRs [4]. Mutations affecting both *SGS1* and *ESC2* cause a synthetic growth defect and elevated gene conversion and joint-molecule formation in *S. cerevisiae* [22]. Moreover, several studies have suggested that the increased genome instability of *mms21* mutants might not be caused by a defect in DNA repair, in contrast to the known repair defects caused by *sgs1* mutations [16–18]. For example, the repair of meiotic DNA DSBs occurs with normal kinetics in *mms21* mutants [23]. In addition, the increased level of SCE in *nsmce2/mms21* mutant mice is not associated with an increase in 53BP1 foci, suggesting a lack of an obvious defect in DNA DSB repair [7]. Together these studies suggest that the genome maintenance functions of the Mms21-Esc2 pathway and Sgs1 might be different.

To gain insight into these questions, we performed a detailed study of the defects caused by the *mms21-CH* mutation, a SUMO E3 ligase-inactive allele of *MMS21*. Our findings show that a diverse array of genome rearrangements accumulate in *mms21-CH* mutants, depending on specific DNA repair pathways available and the nature of genomic sequences involved in the formation of the GCRs observed. Collectively, these findings suggest that spontaneous DNA lesions accumulate in the *mms21-CH* mutant and initiate these genome rearrangements. We further show that Mms21 prevents spontaneous Pol32-dependent break induced replication (BIR), which is also dependent upon the Rrm3 DNA helicase and a subset of the DNA damage checkpoint, but does not involve resolution of recombination intermediates by Sgs1 and does not involve DNA damage-induced sumoylation of Sgs1.

## Results

### Genes in the RAD52 epistasis group are required for the formation of duplication-mediated GCRs in mms21-CH mutant strains

We previously showed that *mms21-CH* caused a substantial accumulation of GCRs in the duplication-mediated GCR or dGCR assay (also called the *yel072w::CAN1/URA3* assay) [4]. In the dGCR assay, non-allelic HR between divergent repetitive sequences on chromosome V and chromosomes IV, X, or XIV resulting in the formation of translocations dominate the selected GCRs in most HR-proficient strains [24,25] (Supplementary Figure 1). In contrast, *mms21-CH* caused only a modest increase in GCR rates in the unique sequence-mediated or uGCR assay (also called the *yel068c::CAN1/URA3* assay) [4], which primarily selects for GCRs mediated by deletions healed by *de novo* telomere additions and various forms of micro- and non-homology mediated translocations [26] (Supplemental Figure 1). To explore this further, we combined the *mms21-CH* mutation with mutations affecting individual genes in the *RAD52* epistasis group in strains containing the dGCR assay or the uGCR assay. We then performed fluctuation analysis to measure the GCR rates of these single and double mutant strains (Figure 1 and Supplementary Table 1).

**Figure 1.**
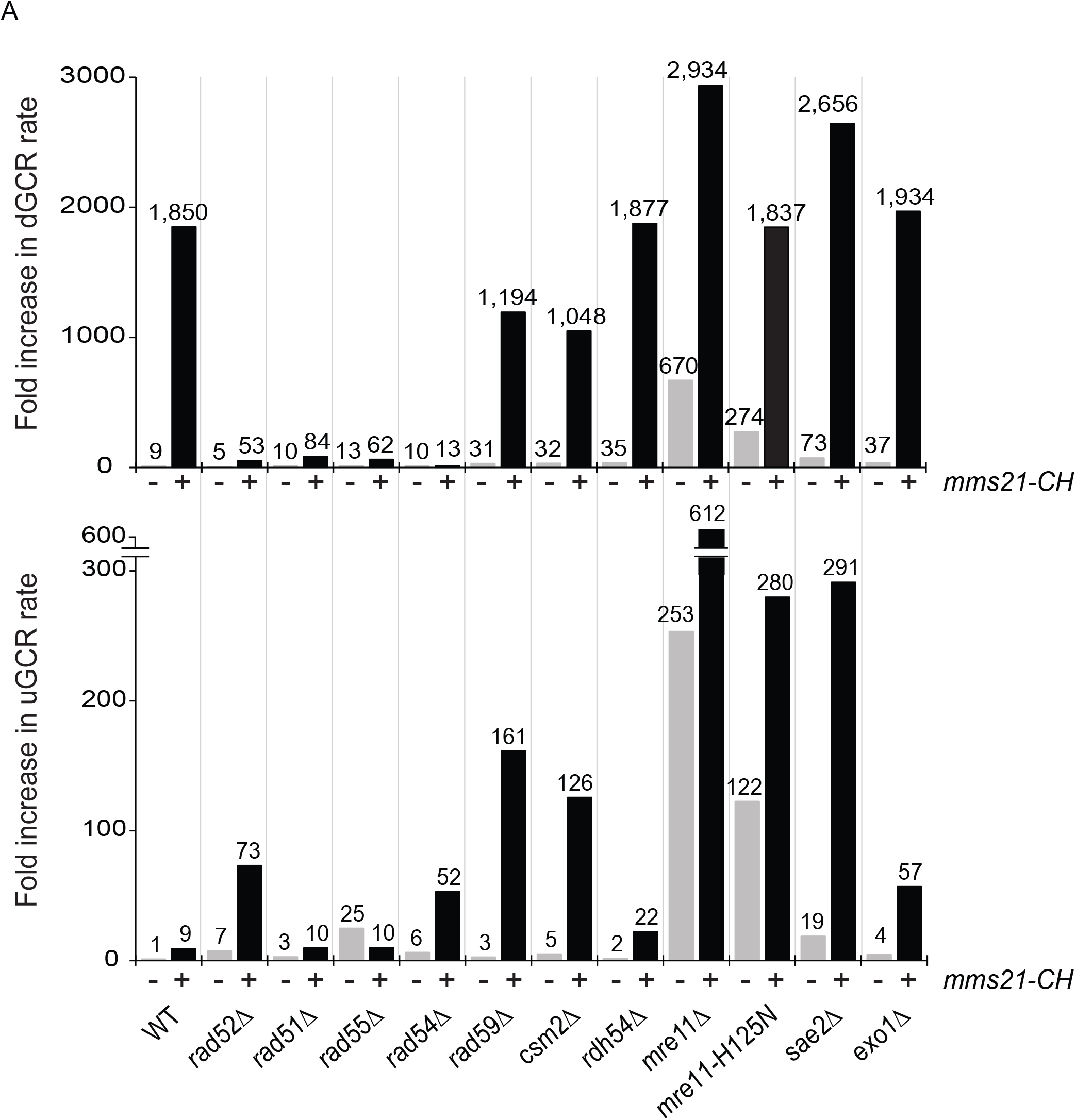
dGCR and uGCR rates caused by mutations of genes in the *RAD52* epistasis group and the *mms21-CH* mutation, a SUMO E3 ligase-null allele. The number above each bar indicates the fold change normalized to the uGCR rate of wild-type strain. Detailed results used to generate the bar graph are shown in Supplementary Table 1.

This analysis of the *RAD52* epistasis group uncovered three main classes of genetic interactions. Class I mutations included deletions in the *RAD51, RAD52, RAD54* and *RAD55* genes required for HR [27]. In this case, deletion of each gene caused a drastic reduction of the increased dGCR rate caused by an *mms21-CH* mutation, indicating a requirement of HR for the formation of GCRs selected in the dGCR assay. Most of the Class I mutations did not cause an increased uGCR rate when combined with an *mms21-CH* mutation, except for *rad52Δ* and *rad54Δ* (as well as a *rad59Δ* mutation; see below); this is consistent with previous observations that HR suppresses GCRs selected in single copy sequence-mediated GCR assays such as the uGCR assay. In contrast, deletion of *RDH54*, which encodes a Rad54 paralog with a role in meiotic HR [28], had little effect on the accumulation of GCRs in the *mms21-CH* mutant.

Class II mutations included deletions of *RAD59* and *CSM2*. Class II mutations partially suppressed the increased GCR rate caused by the *mm21-CH* mutation in the dGCR assay, but caused an increased GCR rate in the uGCR assay when combined with the *mms21-CH* mutation. Rad59 is a stimulatory factor for Rad52 and is important for HR involving shorter repeats or when Rad52 is absent [27]. Csm2 is a subunit of the Shu complex [29], which has been implicated as a regulator of HR, possibly by facilitating the formation of Rad51 filaments [27]; other Shu complex mutations were not tested. Consistent with these accessory roles in HR, deletions of *RAD59* and *CSM2* in the *mms21-CH* mutant modestly reduced the rate of accumulating GCRs in the dGCR assay (Figure 1, upper panel) and substantially increased the rate of accumulating GCRs in the uGCR assay in the *mms21-CH* mutant (Figure 1, lower panel).

Class III mutations included mutations in *MRE11* and *SAE2*. Class III mutations caused at most a modest increase in the increased dGCR rate caused by *mms21-CH*, but caused a substantial increase in the uGCR rate when combined with *mms21-CH*. The Mre11-Rad50-Xrs2 complex, together with Sae2, performs nucleolytic processing of DNA DSBs, leading to 5’-resection at DSBs and an ordered recruitment of HR proteins [27]. Deletion of *MRE11* has been shown to cause substantial increases in the rate of accumulation of GCRs [24], and deletion of *MRE11* in combination with the *mms21-CH* mutation caused an increase in the rate of accumulating GCRs in both the dGCR and uGCR assays compared to the *mms21-CH* single mutant (Figure 1). A mutation inactivating the endonuclease activity of Mre11, *mre11-H125N*, alone caused a 30-fold increase and a 122-fold increase in the rate of accumulating GCRs in the dGCR and uGCR assays, respectively (Figure 1). Interestingly, the *mre11-H125N* mutation did not appreciably affect the dGCR rate of the *mms21-CH* mutant, but caused a further increase in the uGCR rate of the *mms21-CH* mutant, suggesting the involvement of the Mre11 endonuclease activity in suppressing the GCRs selected in the uGCR assay. Sae2 participates in DNA DSB processing by specifically stimulating Mre11 endonuclease activity [30,31]. Like the *mre11-H125N* mutation, deletion of *SAE2* only modestly affected the dGCR rate of the *mms21-CH* mutant, but caused a much larger increase in the uGCR rate of the *mms21-CH* mutant. Thus, the initial nucleolytic processing by Mre11 endonuclease has a critical role in suppressing the formation of the GCRs selected in the uGCR assay in the *mms21-CH* mutant. In contrast, deletion of *EXO1*, which eliminates a key exonuclease that participates in long-range resection of DNA breaks, had very little effect on the rate of accumulating GCRs selected in either the dGCR or uGCR assays in the *mms21-CH* mutant (Figure 1).

### Structures of GCRs formed in the wild-type and the mre11 and mms21-CH mutant strains

To gain further insight into the effects of the loss of *MRE11* and *MMS21* function, we investigated the structures of the GCRs selected in the wild-type, *mms21-CH, mre11Δ, mre11-H125N, mms21-CH mre11Δ*, and *mms21-CH mre11-H125N* mutants. We focused on GCRs selected in the uGCR assay, as the GCRs selected in the dGCR assay are almost exclusively duplication-mediated translocations formed by non-allelic HR between the *DSF1-HXT13* segmental duplication on chromosome V and regions of divergent homology on chromosomes IV, X and XIV, consistent with the HR gene dependency observed for GCRs selected in the dGCR assay in the *mms21-CH* mutant (Supplemental Figure 1). We first characterized the GCRs by testing the individual independent GCR-containing isolates for retention of the telomeric hygromycin resistance marker *hph* located between the telomere and the counter-selectable *CAN1/URA3* cassette on the uGCR assay chromosome and by determining the size of the rearranged chromosome V by Pulse Field Gel Electrophoresis (Figure 2; Supplementary Table 2). GCRs were divided into three groups: rearranged chromosomes that were larger than the wild-type chromosome V (group 1) and chromosomes that were similar to or slightly shorter than the wild-type chromosome V and either lost (group 2) or retained (group 3) the telomeric *hph* marker. We classified GCRs in group 2 as *de novo* telomere addition GCRs, which are formed by the healing of broken chromosomes by the *de novo* addition of a new telomere [32]. *De novo* telomere additions are the predominant form of GCRs selected in uGCR assays in strains without telomerase defects [33–35] and are always associated with loss of the *hph* marker, although it should be noted that rare interstitial deletion GCRs can be associated with deletion of the *hph* marker. Similarly, we classified GCRs in group 3 as interstitial deletion GCRs, in which the deletion is typically associated with non-homology or microhomology breakpoint junctions when selected in GCR assays containing only unique sequences in the breakpoint region like the uGCR assay used here [36].

**Figure 2.**
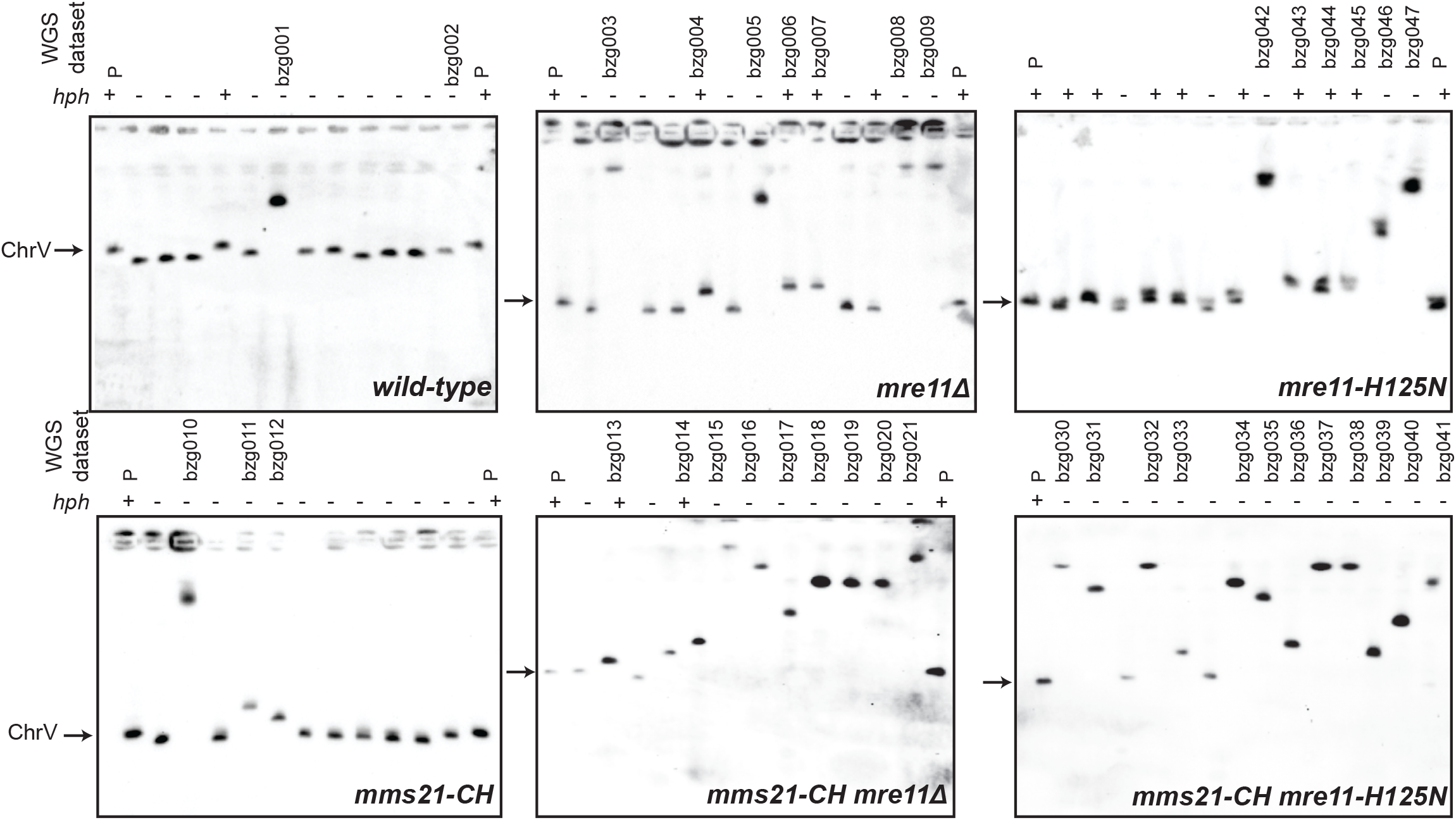
Analysis of the size of the rearranged chromosome V selected in wild-type, *mms21-CH, mre11Δ, mre11-H125N, mms21-CH mre11Δ*, and *mms21-CH mre11-H125N* uGCR strains. Chromosomes were separated by Pulsed-Field Gel Electrophoresis (PFGE) and Southern blotted using a probe for the essential chromosome V gene *MCM3*. The size of chromosome V in the parental strain (P) is indicated by an arrow. The retention (+) or loss (−) of the *hph* marker inserted next to the telomere on the left end of chromosome V is indicated above each lane in the gel. Isolates selected for whole genome sequencing are indicated with their WGS data set name above each lane.

Strains containing GCRs falling into group 1 were subjected to whole genome paired end sequencing to decipher the GCR structures (Supplementary Table 3). In addition to being able to detect all of the mutations and chromosome modifications introduced into the starting strains during strain construction (Supplementary Fig. 2 & 3), we were also able to extensively characterize the structures of the GCR-containing chromosomes (Supplementary Table 4, Supplementary Fig. 4-9). We observed two distinct types of group 1 GCRs: *microhomology-mediated translocations* and *hairpin-mediated inverted duplications*.

In microhomology-mediated translocations, the broken end of a broken chromosome V is fused to another broken chromosome such that the broken chromosome V acquires a fragment of the second broken chromosome that is terminated with a telomere (Figure 3a). Copy number analysis indicated that these fusion events duplicated the non-chromosome V target, and junction sequences revealed only short sequences of identity at the translocation junctions.

**Figure 3.**
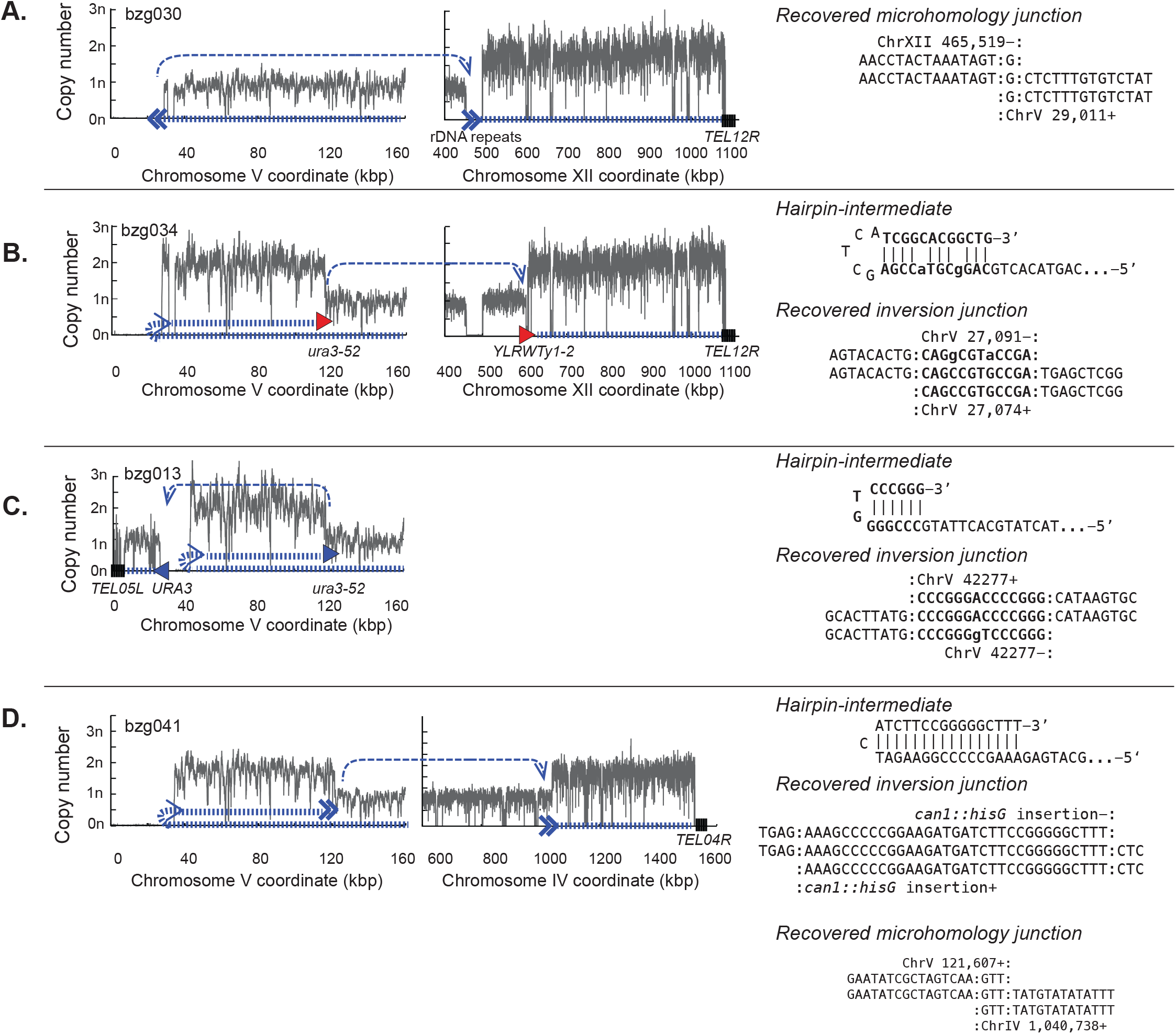
Types of GCRs identified by whole genome sequencing. Analysis of a microhomology-mediated translocation (**A**) and hairpin-mediated inverted duplications that were resolved by homologous recombination between Ty-derived elements (**B**), the homology between *URA3* and *ura3-52* (**C**), and a microhomology-mediated translocation (**D**). **Left**. Copy number analysis of uniquely mapping regions along a portion of the left arm of the assay-containing chromosome V derived from the read depth of uniquely mapping regions of the genome. **Middle**. Copy number analysis that, if present, shows other copy number changes elsewhere in the genome. **Left and Middle**. The path describing the GCR-containing chromosome is illustrated by the hashed thick blue line; the thin dashed blue lines indicate the connectivity between individual fragments that are separated on the reference genome. Homology-mediated translocation junctions are depicted with filled in triangles that point in the direction in which homology element points; junctions involving Ty-related homologies are red and other homologies are blue. Non-homology or micro-homology translocations are shown using two chevrons. Telomeres associated with the GCR (if known) are shown as a series of black vertical lines. **Right**. Sequences of any novel junctions are with the central line in the alignment corresponding to the novel junction. Sequences at the junction that could have been derived from either sequence are surrounded with colons. For GCRs with hairpin-mediated inversions, the inferred structure of the hairpin intermediate is also shown.

In hairpin-mediated inverted duplications the broken end of a broken chromosome V is fused to an inverted copy of itself on the left arm of chromosome V at a position between the *CAN1/URA3* cassette and the first centromeric essential gene (Fig. 3b,c,d). The inversion sites are consistent with a mechanism in which a broken chromosome V is resected to form a 3, overhang that then invades a short stretch of homologous sequence centromeric to the breakpoint followed by replication of the hairpin-terminated chromosome (Supplementary Fig. 10, 11). As previously observed [35], these inverted duplications (also called isoduplications) all underwent additional rounds of rearrangement that either resolved an initial dicentric chromosome or prevented its formation. These secondary rearrangements often, but not always, involved HR between the Ty-or *PAU* gene-related sequences on chromosome V L and a homology elsewhere in the genome (Figure 3b,c; Supplementary Figs. 12-14). Secondary rearrangements that initially appeared to involve HR between *ura3-52* on chromosome V and *YLRCdelta21* on chromosome XII actually proved to target an adjacent full-length Ty element on chromosome XII that was not present in the reference sequence (Supplementary Fig. 15); this full-length Ty element has been previously observed by others [37,38]. A specific secondary rearrangement between a *URA3* fragment in the Ty-inactivated *ura3-52* on chromosome V L and the *URA3* in the *yel068c::CAN1/URA3* cassette first observed in GCRs derived from the *tel1Δ* uGCR strain was also observed here [35]. An additional type of secondary rearrangement was mediated by microhomologies (Fig. 3d); microhomology-mediated secondary rearrangements were not observed in GCRs derived in *tel1Δ* mutants [35]. In most cases, the hairpin-mediated inverted duplications underwent a single secondary rearrangement as described above; however, in a small number of cases multiple rounds of secondary rearrangements were observed leading to the formation of monocentric GCRs (Supplementary Table 4).

### Distribution of GCRs formed in different mutant backgrounds

GCRs selected in the wild-type uGCR strain were primarily *de novo* telomere addition GCRs (Figure 4a; Supplementary Fig. 7), consistent with dominance of *de novo* telomere addition GCRs among the GCRs selected in the “classical” GCR assay [33,39], which lacks large repetitive sequences in the breakpoint region like the uGCR assay used here. In addition, two interstitial deletions and two hairpin-mediated inverted duplications that were resolved by HR between the *ura3-52* allele and the *URA3* gene on the terminal chromosome V telomere-containing fragment were recovered. The spectrum of GCRs obtained from the *mms21-CH* uGCR strain shared this bias towards the formation of *de novo* telomere addition GCRs, with the other GCRs recovered being translocations involving other chromosomes (Figure 4a; Supplementary Fig. 5). In contrast, among the GCRs selected in the *mre11Δ* and *mre11-H125N* single mutant uGCR strains were substantially increased numbers of hairpin-mediated inverted duplications (Figure 4a; Supplemental Fig. 6-7), consistent with role of the Mre11-Rad50-Xrs2 complex in cleaving hairpin structures [31] and subsequent suppression of hairpin-mediated GCRs. In addition, we observed increased numbers of microhomology-mediated translocations in the *mre11Δ* single mutant strain (Supplemental Fig. 5). The *mms21-CH mre11Δ* and *mms21-CH mre11-H125N* double mutants also had increased formation of hairpin-mediated inverted duplications, and the *mms21-CH mre11-H125N* double mutant also had an increased frequency of microhomology-mediated translocations (Figure 4a; Supplementary Fig. 8-9). These biases are also apparent in the GCR rates calculated for each class of GCR (Figure 4b). Remarkably, *MRE11-* deficient strains showed a bias for selection of translocations containing a copy of a long region of chromosome XII R (Figure 4c; Supplementary Fig. 16), which could reflect either a bias due to increased fragility or accessibility of chromosome XII or due to suppression of *mre11-* dependent growth defects by duplication of chromosome XII R. We also observed that 8 of the 10 sequenced *mms21-CH mre11Δ* GCR-containing isolates were disomic for chromosome VIII and 1 of the 10 was disomic for chromosome I (Supplementary Table 2, Supplementary Fig. 17). Taken together, these data are consistent with the idea that the *mms21-CH* mutation increases the total level of DNA damage without substantially biasing the mechanisms involved in forming GCRs, whereas *mre11* defects increase the propensity of damaged DNAs to form hairpin inversions or to undergo microhomology-mediated translocations such as by non-homologous end-joining of two broken chromosomes.

**Figure 4.**
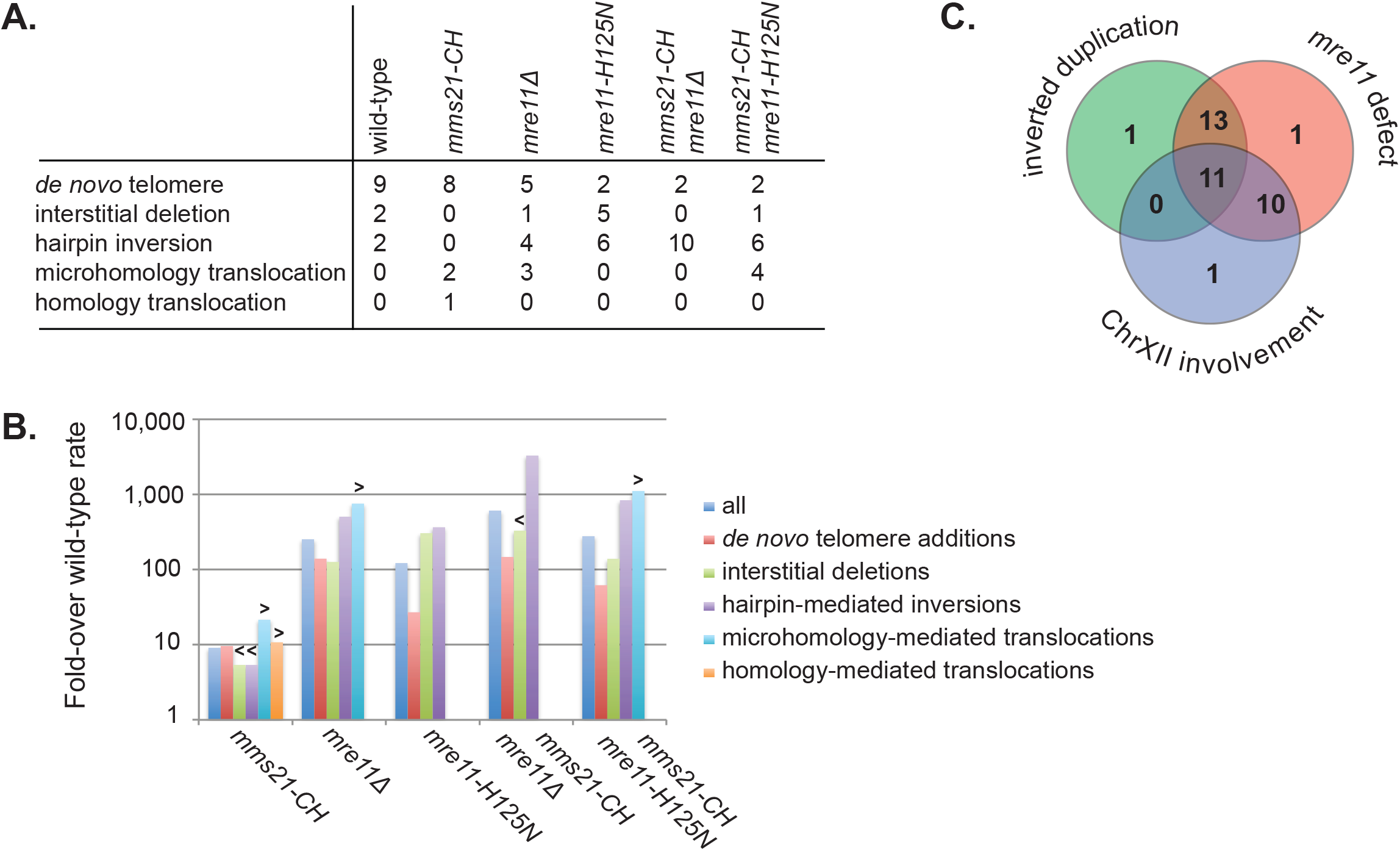
Distribution of GCR types from each mutant analyzed. **A**. Table of the number of each type of rearrangement, *de novo* telomere addition, interstitial deletion, hairpin-mediated inverted duplication, micro- and non-homology-mediated translocations, and homology-mediated translocations. **B**. The fold-increase in the rate of accumulation of all GCRs and each category of GCR as a function of genotype. Less than symbols, “<”, indicate that the fold rate change is less than the maximum possible rate shown (no GCRs of that category were observed in the mutant strain). Greater than symbols, “>“, indicate that the fold rate change is greater than the minimum possible rate shown (no GCRs of that category were observed in the wild-type strain). **C**. Venn diagram of sequenced GCRs demonstrates that strains with *MRE11* defects tend to accumulate GCRs involving inverted duplications and/or chromosome XII.

### Roles of Pol32 and DNA helicases in the accumulation of GCRs in mms21-CH mutant strains

The dramatic HR-dependent increase in the dGCR rate caused by the *mms21-CH* mutation (Figure 1), combined with the fact that *mms21-CH* caused at best modest changes in the spectrum of GCR selected in the uGCR assay (Figure 4 and Supplementary Table 5), suggested that the *mms21-CH* mutation caused an increase in DNA damage without dramatically affecting DNA damage processing. We therefore reasoned that break-induced replication (BIR) might play an important role in GCR formation in *mms21-CH* dGCR strains.

Previous studies of the repair of HO endonuclease-induced DNA DSBs by BIR showed that Pol32, a subunit of DNA polymerases delta and zeta, is required for BIR [40–42]. In contrast, other studies have found that the *pol32Δ* mutation only reduced the efficiency of BIR [43]. We have previously found that a *pol32Δ* mutation does not suppress the wild-type dGCR rate nor does *pol32Δ* eliminate duplication-mediated GCRs [24], suggesting that the role of *POL32* in promoting BIR may be dependent on the nature of the initiating damage. Remarkably, we found that deletion of *POL32* in the *mms21-CH* mutant caused a drastic reduction of the dGCR rate by about 15-fold and a relatively modest increase in its uGCR rate (Figure 5A), consistent with an important role of POL32-dependent BIR in forming dGCRs in the *mms21-CH* mutant.

**Figure 5.**
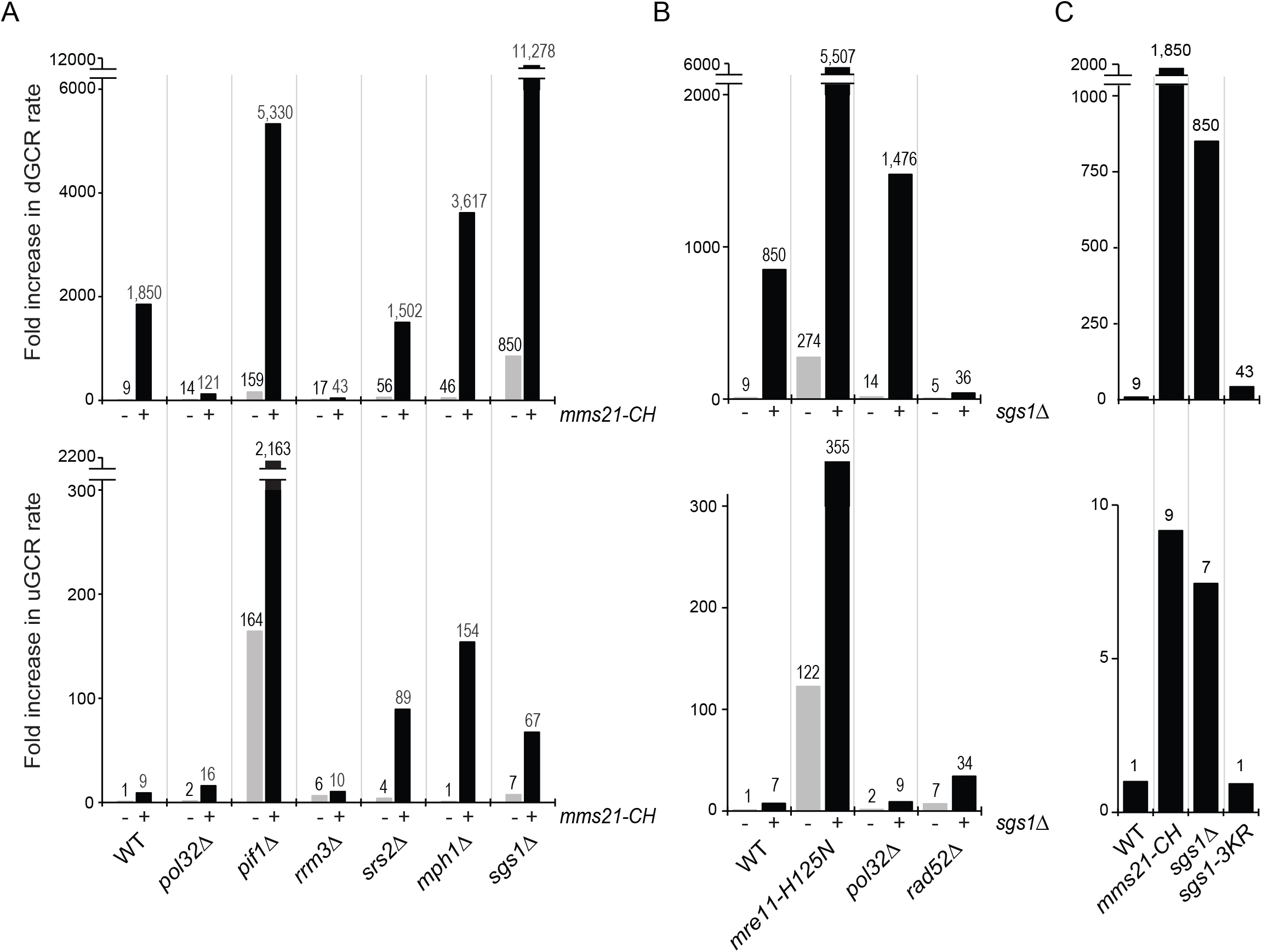
Role of Pol32 and DNA helicases in the formation of GCRs caused by *mms21-CH*. A) dGCR and uGCR rates caused by deleting *POL32, PIF1, RRM3, MPH1, SRS2* and *SGS1* in wild-type and *mms21-CH* mutant strains. B) Effect of combining *sgs1Δ* and *pol32Δ, rad52Δ or mre11-H125N* on the rate of accumulating GCRs. C) Effects of *sgs1-3KR* (K175R, K621R and K831R) mutations on the rate of accumulating GCRs. The number above each bar indicates the fold change normalized to the uGCR rate of wild-type strain. Detailed results used to generate the bar graph are shown in Supplementary Table 5.

Pif1 has been shown to be required for BIR initiated by HO endonuclease-induced DSBs by mediating the DNA synthesis by migration of a D-loop intermediate [40,41]. Pif1 also dissociates telomerase from single-stranded DNA thereby suppressing *de novo* telomere addition GCRs [44]. We found that deleting *PIF1* in the *mms21-CH* mutant caused further increases in both the dGCR and uGCR rates relative to that of the respective single mutants (Figure 5A), which is consistent with the idea that *de novo* telomere addition caused by *pif1Δ* is the pre-dominant mechanism for the formation of GCRs in the *mms21-CH pif1Δ* double mutant [32,35].

We also screened other DNA helicases for their role in forming GCRs in *mms21-CH* mutant strains. *RRM3* encodes a DNA helicase that travels with DNA replication fork [45], but is not known to be required for BIR. Deletion of *RRM3* in the *mms21-CH* mutant caused a drastic and specific reduction (43-fold) in the dGCR rate without appreciably affecting the uGCR rate compared to that of the respective single mutants (Figure 5A), indicating a requirement of Rrm3 in the formation of duplication-mediated GCRs in the *mms21-CH* mutant strains.

The DNA helicase Srs2 acts as an anti-recombinase by disrupting formation of Rad51 filaments [46,47]. In addition, the Smc5/6 complex of which Mms21 is a subunit has been shown to control the recombination activity of the Mph1 helicase [12,48]. Deletion of *SRS2* or *MPH1* in the *mms21-CH* mutant did not appreciably alter the dGCR rate, but caused a drastic increase in the uGCR rate relative to that of the respective single mutants (Figure 5A). This latter result could be explained if Srs2 and Mph1 either suppress hairpin formation and short sequence homology-mediated events that result in secondary rearrangement of some GCRs, or facilitate sister chromatid HR to an extent that suppresses GCRs selected in the uGCR assay, but not those selected in the dGCR assay.

The Sgs1 helicase has a major role in specifically suppressing dGCRs [15,24], and this has been attributed to its role in preventing crossovers during the resolution of HR intermediates [49]. Interestingly, combining an *sgs1Δ* with the *mms21-CH* mutation resulted in synergistic increases in both dGCR and uGCR rates relative to the respective single mutants (Figure 5A), indicating Mms21 and Sgs1 function in distinct pathways to prevent the formation of GCRs. To explore this further, we analyzed the effects of mutating *RAD52, MRE11* and *POL32* in the *sgs1Δ* mutant. Deletion of *RAD52* and the *mre11-H125N* mutation caused similar effects in *sgs1Δ* and *mms21-CH* mutants (Figures 1 and 5). In contrast, deletion of *POL32* caused an increase in the dGCR rate of the *sgs1Δ* mutant whereas deletion of *POL32* in the *mms21-CH* mutant reduced the dGCR rate more than 10-fold (Fig. 5B). Thus, the formation of duplication-mediated GCRs in the *mms21-CH* and *sgs1Δ* mutants had distinctly different requirements for Pol32. Because deletion of *RRM3* is lethal in an *sgs1Δ* mutant [50], we could not compare the roles of Rrm3 in the *mms21-CH* and *sgs1Δ* mutants.

Recent studies showed that Mms21 specifically catalyzes sumoylation of Sgs1 in response to treatments with DNA alkylating agents [10,19,20]. We found that the *sgs1-3KR* mutation that eliminates the sumoylation sites on Sgs1 did not cause a comparable increase in GCR rates like that seen in *sgs1Δ* mutants (Figure 5C). Although we cannot exclude the possibility that a low and undetectable level of Sgs1 sumoylation occurs in the *sgs1-3KR* mutant, this result indicates that the major DNA damage-induced sumoylation of Sgs1 does not appear to have an appreciable role in preventing spontaneous GCRs.

### Mutation affecting MMS21 induces spontaneous DNA lesion and activates DNA damage checkpoint to promote dGCRs

The above findings are consistent with the idea that *mms21-CH* causes accumulation of spontaneous DNA lesions that underlie the formation of a diverse range of GCRs. Cells have evolved a signal transduction pathway, the DNA damage checkpoint, to detect endogenous DNA lesions [51]. We therefore examined the Rad53 kinase, which becomes hyperphosphorylated in the presence of such DNA damage and migrates with a slower electrophoretic mobility than non-phosphorylated Rad53. Treatment by the DNA alkylating agent methyl methane sulfonate (MMS) caused a pronounced electrophoretic mobility shift of fully activated Rad53 to slower migrating species (Figure 6A). An increased but sub-stoichiometric amount of Rad53 was found to show slower gel mobility in an *mms21-CH* mutant that was not treated with MMS compared to untreated wild-type, indicating that Rad53 is partially activated in *mms21-CH* mutants. Deletion of *RAD9*, which encodes an adaptor protein that acts to promote DNA damage-induced activation of Rad53, reduced the amount of the slower migrating species of Rad53 to an undetectable level in the *mms21-CH* mutant that was not treated with MMS; note that MMS-induced Rad53 phosphorylation still occurs in a *rad9Δ* mutant due to the redundant role of Mrc1 in mediating Rad53 activation [52–54]. Ddc2, together with the Mec1 kinase, is recruited to RPA-coated single stranded DNA at the sites of DNA damage where it can be visualized as sub-nuclear foci [55]. A higher incidence of Ddc2 foci was seen in the untreated *mms21-CH* mutant compared to untreated wild-type cells (Figure 6B). Together, these results suggest that elevated levels of endogenous DNA lesions occur in the *mms21-CH* mutant.

**Figure 6.**
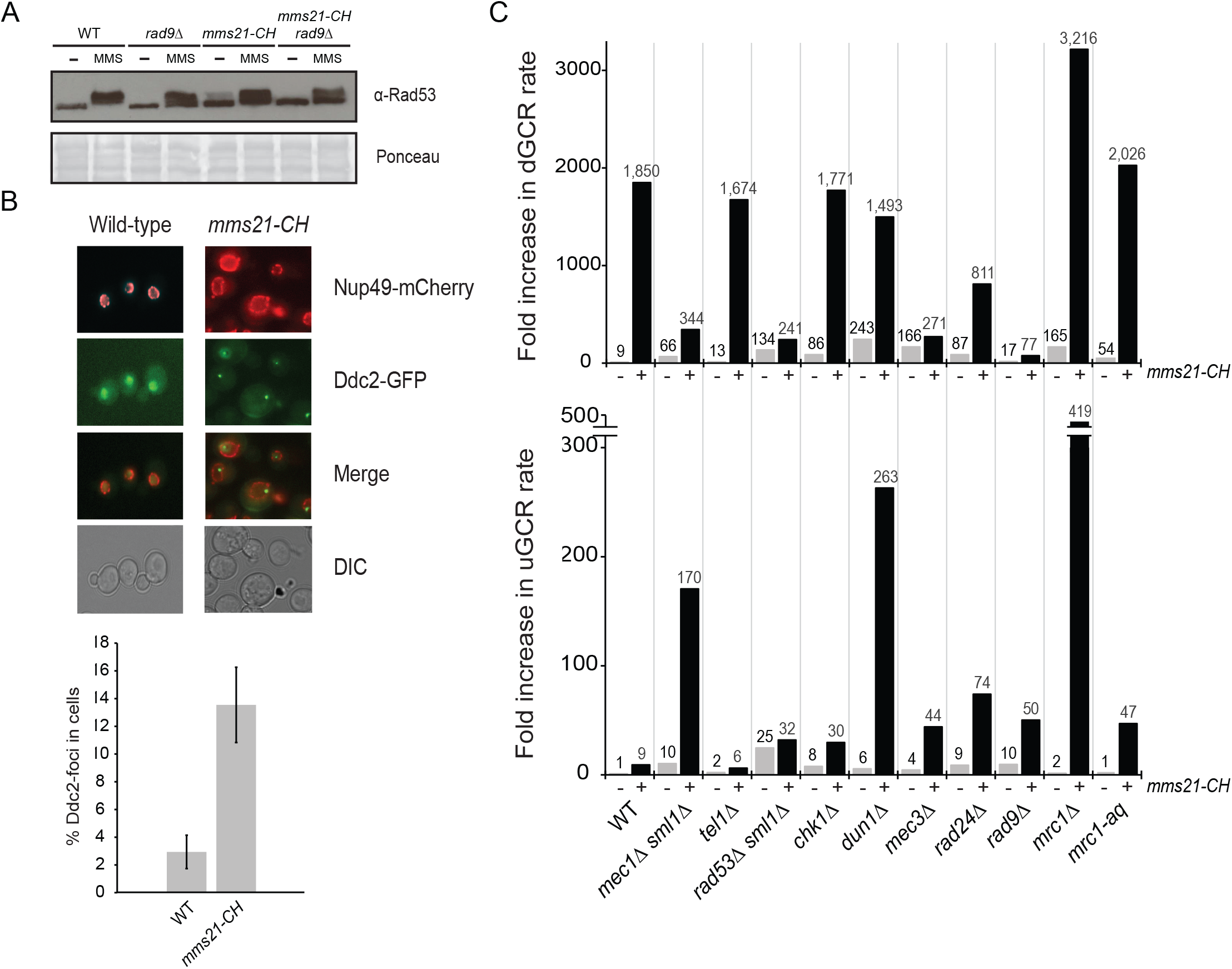
Role of DNA damage checkpoint in the formation of GCRs. A) Rad53 gel shift assay to examine Rad53 activation in WT, *mms21-CH, rad9Δ* and *rad9Δ mms21-CH* mutants. B) Spontaneous Ddc2 foci in *mms21-CH* mutant and WT, which also contain Nup49-mCherry to mark the nuclear envelope. Bar graph indicates the percentage of Ddc2-foci within the nuclear envelope (marked by Nup49-mCherry). Error bars represent the standard deviation from three replicate experiments using two biological isolates per strain per replicate. C) dGCR and uGCR rates caused by mutations of DNA damage checkpoint genes with or without *mms21-CH*. The number above each bar indicates the fold change normalized to the uGCR rate of wild-type strain. Detailed results used to generate the bar graph are shown in Supplementary Table 6.

We next asked whether the DNA damage checkpoint influences the formation of GCRs in the *mms21-CH* mutant (Figure 6C and Supplementary Table 6). The DNA damage checkpoint involves two partially redundant protein kinases, Mec1 and Tel1. While Mec1 has a major role in controlling the DNA damage response, Tel1 has an important role in telomere length maintenance in wild-type cells and in checkpoint responses in *mec1* mutants [56,57]. We found that deletion of *MEC1* caused a 5-fold reduction in the dGCR rate of the *mms21-CH* mutant (Figure 6C, upper panel) and a substantial increase in the uGCR rate of the *mms21-CH* mutant (Figure 6C, lower panel). Unlike the *mec1Δ* mutation, deletion of *TEL1* caused little of no change in the dGCR and uGCR rates of the *mms21-CH* mutant. Like the deletion of *MEC1*, deletions of the *MEC3, RAD24* and *RAD9* genes involved in the DNA damage checkpoint, caused varying degrees of reduction of the dGCR rate of the *mms21-CH* mutant with the *rad9Δ* mutation causing the greatest reduction (~ 24-fold). In contrast, deletions of the *MEC3, RAD24* and *RAD9* genes in the *mms21-CH* mutant resulted in increases in the uGCR rate, consistent with the known role of DNA damage checkpoint in suppressing GCRs mediated by single-copy sequences [58].

Unlike deletion of *RAD9*, deletion of *MRC1* caused a modest (2-fold) increase in the dGCR rate of the *mms21-CH* mutant (Figure 6C, upper panel). Interestingly, deletion of *MRC1* caused a drastic increase in the uGCR rate of the *mms21-CH* mutant (Figure 6C, lower panel). Because Mrc1 also has a role in DNA replication [59], we next examined the *mrc1-AQ* mutant, all of whose Mec1 consensus phosphorylation sites are mutated to non-phosphorylatable alanines and is thus unable to mediate Rad53 activation [60]. We found that the *mrc1-aq* mutation did not appreciably alter the dGCR rate of the *mms21-CH* mutant although it did cause an increase in the uGCR rate of the *mms21-CH* mutant, but not to the extent seen with the *mrc1Δ* mutation. Rad53, Chk1, and Dun1 are the downstream effector kinases of the checkpoint pathways. Deletion of *RAD53* reduced the dGCR rate of the *mms21-CH* mutant by about 9-fold (Figure 6C, upper panel), while deletion of *CHK1* did not appreciably alter the dGCR rate of the *mms21-CH* mutant and caused a small increase in the uGCR rate of the *mms21-CH* mutant. Although deletion of *DUN1* did not appreciably alter the dGCR rate of the *mms21-CH* mutant, it caused a synergistic increase in the uGCR rate of the *mms21-CH* mutant (Figure 6C, lower panel). These data support the idea that the Mec3/Rad24-Rad9-Mec1-Rad53 pathway plays a role in promoting the GCRs selected in the dGCR assay in the *mms21-CH* mutant while suppressing the GCRs selected in the uGCR assay in the *mms21-CH* mutant.

## Discussion

Mutations affecting the Mms21 SUMO E3 ligase cause substantially increased accumulation of DNA damage intermediates and increased accumulation of GCRs [4,5,10]. Here we have shown that the Mms21 E3 ligase plays an important role in suppressing the formation of GCRs selected in the dGCR assay, which are typically translocations mediated by non-allelic HR (Figure 7). The duplication-mediated GCRs formed in Mms21 E3 ligase mutants appear to be formed by *POL32-* dependent BIR in contrast to those formed in wild-type strains, which have little dependence on Pol32 [24]. The increased rate of accumulating GCRs selected in the uGCR assay caused by *mms21-CH* mutations, which does not reflect the formation of duplication-mediated GCRs, is not accompanied by a change in the spectrum of GCRs relative to the spectrum of GCRs selected in the uGCR assay in wild-type strains. This suggests that the *mms21-CH* mutation causes increased levels of DNA damage that trigger the DNA damage checkpoint and are processed into GCRs. We cannot rule out the possibility that Mms21 plays roles in some DNA repair pathways; however, the accumulated evidence presented here suggests that Mms21 suppresses genome instability primarily by preventing the formation of some initiating DNA damage. One possibility is that Mms21 promotes sister chromatid allelic HR through its role in the Smc5/Smc6 complex and this reduces the level of DNA damage that would otherwise result in GCRs (Figure 7). In this regard, it is intriguing that Mms21, along with its positive regulator Esc2, promotes sumoylation of a number of targets including cohesin and condensin subunits [4,61–63].

**Figure 7.**
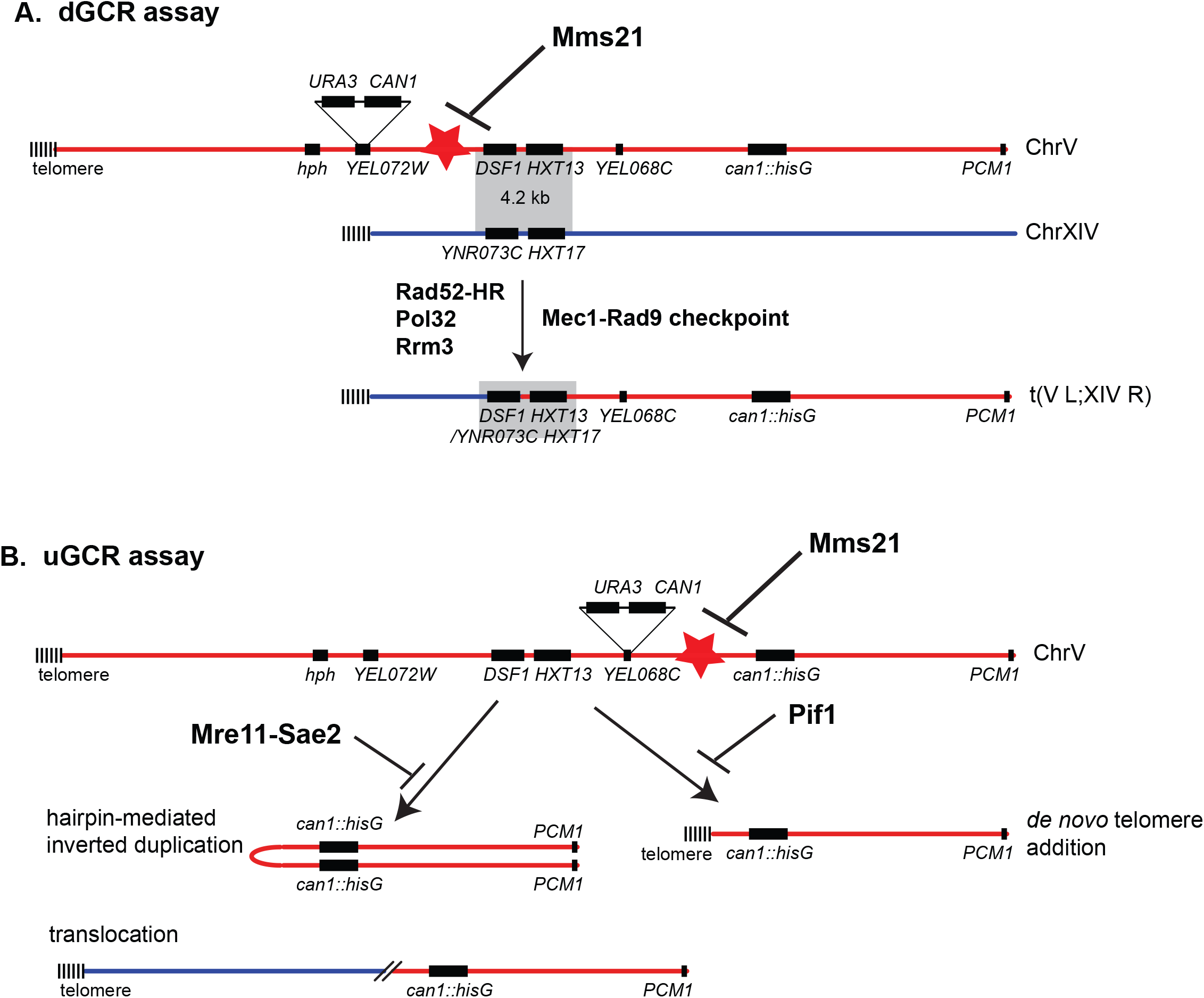
Summary of the functions of the Mms21 E3 ligase pathway in preventing genome rearrangements. A) dGCR assay detects BIR events, driven by Pol32, Rrm3, DNA damage checkpoint and Rad52 dependent non-allelic HR. B) uGCR assay detects micro-homology mediated translocations and hairpin mediated inverted duplications prevented by the Mre11-Sae2 endonuclease, and *de novo* telomere addition prevented by the Pif1 helicase. These GCR events are stimulated by inactivating Mms21 E3 ligase to cause DNA lesions, depicted as red stars.

The GCRs selected in the dGCR assay are primarily translocations formed by non-allelic HR between the *DSF1-HXT13* region on chromosome V and divergent homologous sequences elsewhere in the genome [24]. The genetic requirements for the duplication-mediated GCRs that are formed at increased rates in *mms21-CH* mutants are consistent with the idea that these GCRs are formed by non-allelic HR mediated by BIR; the increased dGCR rates were greatly reduced when the genes required for HR were deleted, were partially reduced when accessory HR genes were deleted and were greatly reduced when the *POL32* gene required for BIR initiated by HO endonuclease-induced DSBs was deleted [42]. The fact that the *pol32Δ* mutation does not decrease the dGCR rate in wild-type cells [24] may suggest that the DNA lesions that initiate the formation of GCRs in *mms21-CH* mutants are more similar to HO endonuclease-induced DSBs or readily converted to such DSBs than the DNA lesions that underlie duplication-mediated GCRs in wild-type cells. This possibility is also consistent with the constitutive activation of the DNA damage checkpoint in *mms21-CH* mutants as indicated by increased Rad53 hyperphosphorylation and increased levels of Ddc2 foci in *mms21-CH* mutants (Figure 6). The Pif1 DNA helicase has also been shown to be required for BIR initiated from HO endonuclease-induced DSBs [40,41]; however, a *pif1Δ* mutation did not decrease the dGCR rate of *mms21-CH* mutants, although the predicted effect of the *pif1Δ* mutation on BIR could be masked by the large increase in the rate of *de novo* telomere addition GCRs in *pif1Δ* strains [32,36]. Moreover, the increase in the *mms21-CH* rates in the uGCR and dGCR assays caused by a *pif1Δ* mutation is consistent with the possibility that *MMS21* suppresses the formation of DNA damage; loss of the Pif1 DNA helicase causes increased GCR rates when combined with many different mutations that lead to increased levels of DNA damage. Together, these data argue that the increased dGCR rate seen in *mms21-CH* mutants is the result of increased non-allelic HR that is most likely mediated by BIR due to increased levels of DNA lesions that are substrates for BIR. We have not ruled out the possibility that the *mms21-CH* mutation also alters the activity of some of the HR proteins.

We have found that duplication-mediated GCRs that occur at increased rates in *mms21-CH* mutants depend on both the Rrm3 DNA helicase and the DNA damage checkpoint. Unlike Pif1, Rrm3 is not known to be required for HO-induced DSB-mediated BIR, and purified Rrm3 is not able to replace Pif1 in the extension of D-loops by DNA polymerase delta *in vitro* [41]. Why Rrm3 is required for the formation of duplication-mediated GCRs in *mms21-CH* mutants is not clear. Rrm3 might act to generate GCR-inducing DNA lesions in *mms21-CH* mutants; one possible mechanism could involve *mms21-CH*-induced aberrant processing of replication forks stalled at protein-DNA complexes [45] when Mms21 targets such as cohesin subunits are sumoylated. Alternatively, Rrm3, a homolog of Pif1 [64], might also play a role in promoting the formation of mms21-CH-induced GCRs mediated by BIR similar to how Pif1 acts [40,41]. We have also identified the DNA damage checkpoint as promoting the formation of duplication-mediated GCRs in *mms21-CH* mutants. The DNA damage checkpoint has well-documented roles in promoting the homology search during HR [65] which could promote the production of the GCRs selected in the dGCR assay. Remarkably, deletion of *RAD9* caused a specific reduction of the dGCR rate, consistent with the formation HR-initiating DNA lesions that are not associated with active or stalled replication forks, but rather might be DSBs or some other type of DNA damage that is converted to DSBs.

Loss of either Mms21/NSMCE2 or Sgs1/BLM causes increased levels of aberrant HR intermediates, SCE and GCRs [4,5,10,11,13], and sumoylation of Sgs1 by Mms21 prevents the formation of aberrant HR intermediates response to alkylation damage [19,20]. Despite these similarities, the increase in the dGCR and uGCR rates caused by combining the *mms21-CH* and *sgs1Δ* mutations argues strongly that these proteins act in different pathways that suppress the formation of GCRs. Consistent with this conclusion, deletion of *POL32* had differing effects on dGCR rates in *mms21-CH* strains relative to *sgs1Δ* strains. Sgs1 is important in resolving HR intermediates [16]; hence, the GCR-based genetic interactions between *sgs1Δ* and *mms21-CH* mutations seen here suggest that Mms21 prevents the formation of damage that underlies aberrant HR and that Sgs1 acts to edit these aberrant HR intermediates to prevent non-allelic HR.

The increased accumulation of DSBs or damage that can be converted to DSBs in *mms21-CH* mutant strains is also consistent with the structures of the GCRs selected in the uGCR assay as determined by whole genome sequencing. The *mms21-CH* single mutants have increased rates of accumulating *de novo* telomere addition GCRs and microhomology-mediated translocation GCRs, which reflect different mechanisms of healing broken chromosomes. Interestingly, mutations affecting *MRE11* caused a large increase in hairpin-mediated inverted duplications as well as microhomology-mediated translocations when combined with an *mms21-CH* mutation. These results are consistent with increased formation of DSBs in an *mms21-CH* mutant combined with the inability of *mre11* mutants to cleave DNA hairpins generated from these DSBs [31] and to prevent microhomology-mediated translocations at these DSBs [33].

Together, the findings presented here argue that mutations inactivating the Mms21 E3 ligase lead to an accumulation of DNA lesions that are either DNA DSBs or easily convertible to DNA DSBs, and that these lead to diverse genome rearrangements, depending on the available DNA repair pathways (Figure 7). Considering Mms21 has been shown to sumoylate proteins with essential roles in DNA replication and chromosome maintenance, including the MCM2-7 replicative helicase and SMC complexes [4,6], it is tempting to speculate that defects in Mms21 sumoylation could cause DNA DSBs and modulate their repair especially via non-allelic HR pathways.

## Acknowledgements

We would like to thank Dr. Marco Foiani for anti-Rad53 antibody, and members of the Zhou and Kolodner labs for discussion during the course of this study.

## Material and methods

### S. cerevisiae strain construction and genetic methods

Standard *S. cerevisiae* genetics method was used to introduce mutations. *S. cerevisiae* strains used in this study are listed in Supplementary Table 7. Generation of the *sgs1-3KR* mutation involved two steps by first deleting the N-terminal region of Sgs1 from −425bp to 2600bp and then repairing it using PCR products containing the *sgs1-3KR (K175R, K621R and K831R)* mutations with a *HIS3* marker located at 435bp upstream of the starting codon of Sgs1 to preserve its native promoter. DNA sequencing was used to confirm the integration of *sgs1* mutations. Methods used for fluctuation analysis to determine GCR rates have been described previously [66].

## Microscopy

Cells were grown in CSM medium to log phase and examined by live imaging using Olympus BX43 fluorescence microscope with a 60x, 1.42 PlanApo N Olympus Oil immersion objective. GFP and mCherry fluorescence were detected using a Chroma FITC filter set and a TxRed filter set respectively and captured with a Qimaging QIClick CCD camera. Images were captured using Meta Morph Advanced 7.7 imaging software. 200-400 cells were imaged and counted for each experiment. Figures were prepared in Adobe Photoshop, keeping processing parameters constant within each experiment.

## PFGE and Southern blotting

DNA plugs for PFGE were prepared as described [67]. Electrophoresis was performed using a Bio-Rad CHEF-DRII apparatus at 6 V/cm, with a 60 to 120 s switch time for 25 h. The gels were stained with ethidium bromide and imaged. The DNA in the gel was transferred to Hybond-XL membranes by neutral capillary blotting. The DNA was crosslinked to the membrane by UV irradiation in a Stratalinker^™^ (Stratagene) apparatus at maximum output for 60 seconds. The *MCM3* probe was generated by amplifying *MCM3* from genomic DNA using the primers 5′-CTGTGCAAGAAATGCCCGAAATG-3′ and 5′-GCCCCGGAGTTGGAATGCTC-3′ followed by random primer labeling of the PCR product with the Biotin DecaLabel DNA Labeling Kit (Thermo Scientific). Probe hybridization was performed at 50°C for 1 hr. Biotin signal was detected using Chemiluminescent Nucleic Acid Detection Module Kit (Thermo Scientific).

## Whole genome paired-end sequencing

Multiplexed paired-end libraries were constructed from 2 μg of genomic DNA purified using the Purgene kit (Qiagen). The genomic DNA was sheared using M220 focused-ultrasonicator (Covaris) and end-repaired using the End-it DNA End-repair kit (Epicentre Technologies). Common adaptors from the Multiplexing Sample Preparation Oligo Kit (Illumina) were then ligated to the genomic DNA fragments, and the fragments were then subjected to 18 cycles of amplification using the Library Amplification Readymix (KAPA Biosystems). The amplified products were fractionated on an agarose gel to select 600 bp fragments, which were subsequently sequenced on an Illumina HiSeq 4000 using the Illumina GAII sequencing procedure for paired-end short read sequencing. Reads from each read pair were mapped separately by bowtie version 2.2.1 [68] to a reference sequence that contained revision 64 of the *S. cerevisiae* S288c genome (http://www.yeastgenome.org), *hisG* from *Samonella enterica*, and the *hphMX4* marker. Sequence data is available from National Center for Biotechnology Information Sequence Read Archive under accession number: SRP106876.

## Rearrangement and copy number analysis of paired-end sequencing data

Chromosomal rearrangements were identified after bowtie mapping by version 0.6 of the Pyrus suite (http://www.sourceforge.net/p/pyrus-seq) [35]. Briefly, after removal of PCR duplicates, read pairs in which both reads uniquely mapped were used to generate the read depth and span depth copy number distributions. The read depth copy number distribution is the number of times each base pair was read in a sample; read depth distributions were the distributions plotted to examine copy number (Supplementary Figures 4-9) as this distribution is less distorted than the span depth distribution in regions adjacent to repetitive elements. The span depth copy number distribution is the number of times each base pair in a sample was contained in a read or spanned by a pair of reads; span depth distributions were used to statistically distinguish real rearrangements identified by junction-defining discordant read pairs from discordant read pairs that were noise in the data. Read pair data were then analyzed for junction-defining discordant read pairs that indicated the presence of structural rearrangements relative to the reference genome. Identified rearrangements included junctions produced during strain construction, such as the *his3Δ200* deletion (see Supplementary Figures 2-3), or GCR-related rearrangements (see Supplementary Figures 4-9). Associated junction-sequencing reads, which were reads that did not map to the reference but were in read pairs in which one end was adjacent to discordant reads defining a junction, were used to sequence novel junctions. Most hairpin-generated junctions (Supplementary Figure 11) could be determined using alignments of junction-sequencing reads. For junctions formed by HR between short repetitive elements (Supplementary Figures 12-14) and for problematic hairpin-generated junctions (Supplementary Figure 11), the junction sequence could be derived by alignment of all reads in read pairs where one read was present in an “anchor” region adjacent to the junction of interest and the other read fell within the junction to be sequenced. Similar strategies involving the alignment of reads paired with reads present in “anchor” regions also were used to sequence *de novo* telomere addition junctions (Supplementary Figures 4-9) and to identify the *“YLRWTy1-4”* Ty element that was not present in the reference genome (Supplementary Figure 15).

### Rad53 gel shift analysis

Protein extracts for Western blot analysis was prepared using a TCA (trichloroacetic acid) extraction. To examine Rad53 electrophoretic mobility we used an anti-Rad53 monoclonal antibody (EL7E1 serum) from mouse, a gift from Dr. Marco Foiani.

